# Trunk stiffness of the American giant millipede during terrain bridging

**DOI:** 10.64898/2026.11.11.765885

**Authors:** Alyssa Micheli, Timothy Perez, Colleen Unsworth, Apoorva Khairnar, Henry Astley, Noel Naughton

## Abstract

Slender, flexible bodies allow animals such as millipedes to regularly traverse heterogeneous, uneven terrain and complex obstacles by using their body’s compliance to conform to the terrain. Orchestrating a slender body’s many degrees of freedom poses a control challenge, with possible strategies ranging in complexity and cost, from complex muscle actuation patterns to simple passive compliance. Here we investigate these behavioral strategies by examining American giant millipedes (*Narceus americanus*) bridging between terrains of different heights. We model millipede trunk mechanics using Cosserat rods and compare resulting equilibrium configurations with experimental data to determine bending stiffnesses that best recreates their posture and test for regional and temporal variations. We find that a highly compliant, passive rod best captures millipede poses during periods where its middle section is unsupported. This supports the hypothesis that millipedes do not strongly actively stiffen their trunk during terrain bridging, relying instead on their body’s passive compliance.

## INTRODUCTION

Long, slender bodies are widespread among animals, and the flexibility of these bodies allows animals to conform to and engage with structurally complex terrain [1–3]. This flexibility arises from either a high number of articulated joints (e.g. snakes, millipedes) [3–5] or a continuously deformable structure (e.g. elephant trunks, worms, octopus arms) [6–8] and, consequently, requires control of an extremely large or effectively infinite number of degrees of freedom. This allows a wide range of control strategies, ranging from finely controlled muscle activity which varies both spatially and temporally[9–12] to reliance of simple passive body stiffness[13, 14]. These strategies have costs, both metabolically and computationally, but can also be constrained by the anatomy and neural control mechanisms of a given species.

Millipedes (Arthropoda, Myriapoda, Diplopoda) [15] use their long, flexible bodies to conform to complex, restrictive terrain and thereby maintain contact with the uneven ground [3]. This ground contact is necessary as millipede legs lack adhesive properties and many species, including *Narceus americanus*, lack claws, and thus millipedes rely on leg contact for support, resulting in poor climbing compared to other arthropods [16]. This body flexibility is achieved via a series of repeating segments of legs, ganglia, and muscles. This series of segments is comprised of both rigid sclerites and softer connecting material, with muscles often anchored to the sclerites at one or both ends [3, 4]. Sclerites are connected through a ball-and-socket joint which provides rotation in three degrees of freedom [3, 17], allowing for the bending and twisting observed in millipedes. Millipede trunk muscles are limited to connecting adjacent segments [17]. Inves-tigations on their limb movement patterns suggest a decentralized pattern of neural control [18, 19], which likely extends to overall truck control.

Previous investigations of myriapoda locomotion have examined the dynamic gait cycle of millipedes and centipedes across both flat and uneven terrain, generally focusing on leg gait independent of trunk pose in biological [20, 21] and robophysical [22–25] contexts. Some work has also considered trunk pose, in particular its relation to leg activation during the gait cycle [26–29] and its role in behaviors such as horizontal movement [16] and curling [30]. However, millipede trunk control during terrain changes and unsupported locomotion remains less understood. In supported terrain bridging, such as transitions in continuous slopes, the millipede maintains contact and conforms to the shape of the terrain. In contrast, during unsupported terrain bridging, such as crossing steps of different heights, the middle section of the millipede looses contact with the terrain, leading to its pose becoming heavily influenced by the the millipede’s trunk’s material properties, such as its bending stiffness, which may be passive or may be actively modulated by muscle activity. Such muscle activity could be uniform, or could vary along the body and/or over time.

Here, we investigate the bending stiffness of *N. americanus* as it traverses a vertical step in terrain. During this action, legs in its middle section lose contact with the ground and the millipede’s trunk becomes suspended off the ground. We develop a simplified model of millipede mechanics by considering the millipede as a passive beam using a Cosserat rod model. By varying the stiffness of the beam, we can test whether the millipede actively stiffens the trunk. Furthermore, closer examination of any discrepancies between the millipede behavior and the uniform beam model would detect regional stiffening patterns along the body, or differences in stiffness over time.

## MATERIALS AND METHODS

### Millipede gap crossing experiments

We purchased 10 American giant millipedes (*Narceus americanus*) from a commercial dealer, and maintained them in a container of topsoil and leaf litter, with assorted plant material as food. Experimental terrain was constructed from hand-cut closed-cell rigid house insulation foam and covered with masking tape for traction and uniform friction. Ledge overhangs were used to induce traversal with a suspended segment, as millipedes would simply scale vertical ledges with legs maintaining continuous contact. Terrain depth was slightly more than one body width and sandwiched between white posterboard in the back and clear acrylic in front to restrict movement to being parallel to the camera’s imaging plane. All trials were recorded with a Nikon D3500 camera at 1080p resolution and 60 frames per second at room temperature. A clear ruler was included to provide scale. Trials consisted of millipedes traversing steps at heights ranging from 1.0 to 2.5cm and overhang angles of 22.5◦ and 45◦. 32 trials total were conducted, with 16 taken at a low position, 9 at a high position, and 7 at a medium position. Within these trials the amount of ledge overhang and direction of travel were also varied, with 13 videos at 45 degrees up-wards, 12 videos at 22.5 degrees upwards, and 7 videos at 22.5 degrees downwards, though this was not found to have a major impact of millipede behavior. Frames from videos were extracted every 10 frames leading to a timestep between frames of 0.167 seconds. This produced a total of 325 frames at the low position, 290 frames at the high position, and 212 frames at the medium position.

Video processing. To extract millipede poses, for each frame, millipede bodies were isolated from the background environment using a custom-trained YOLO11 deep learning instance segmentation model [31, 32] implemented in Python. Training data was generated by manually labeling millipede outlines across a range of trials using Roboflow [33]. This binary segmentation mask was then shrunk using a morphological thinning operation to produce a midline of the millipede’s pose to which a smoothing spline, sampled at fixed 10 pixel arc-length intervals, was fit (Fig. 1a). The video’s ruler markings were isolated using Hough Line transforms [34] and processed to produce a pixel-to-millimeter scale.

**FIG 1.**
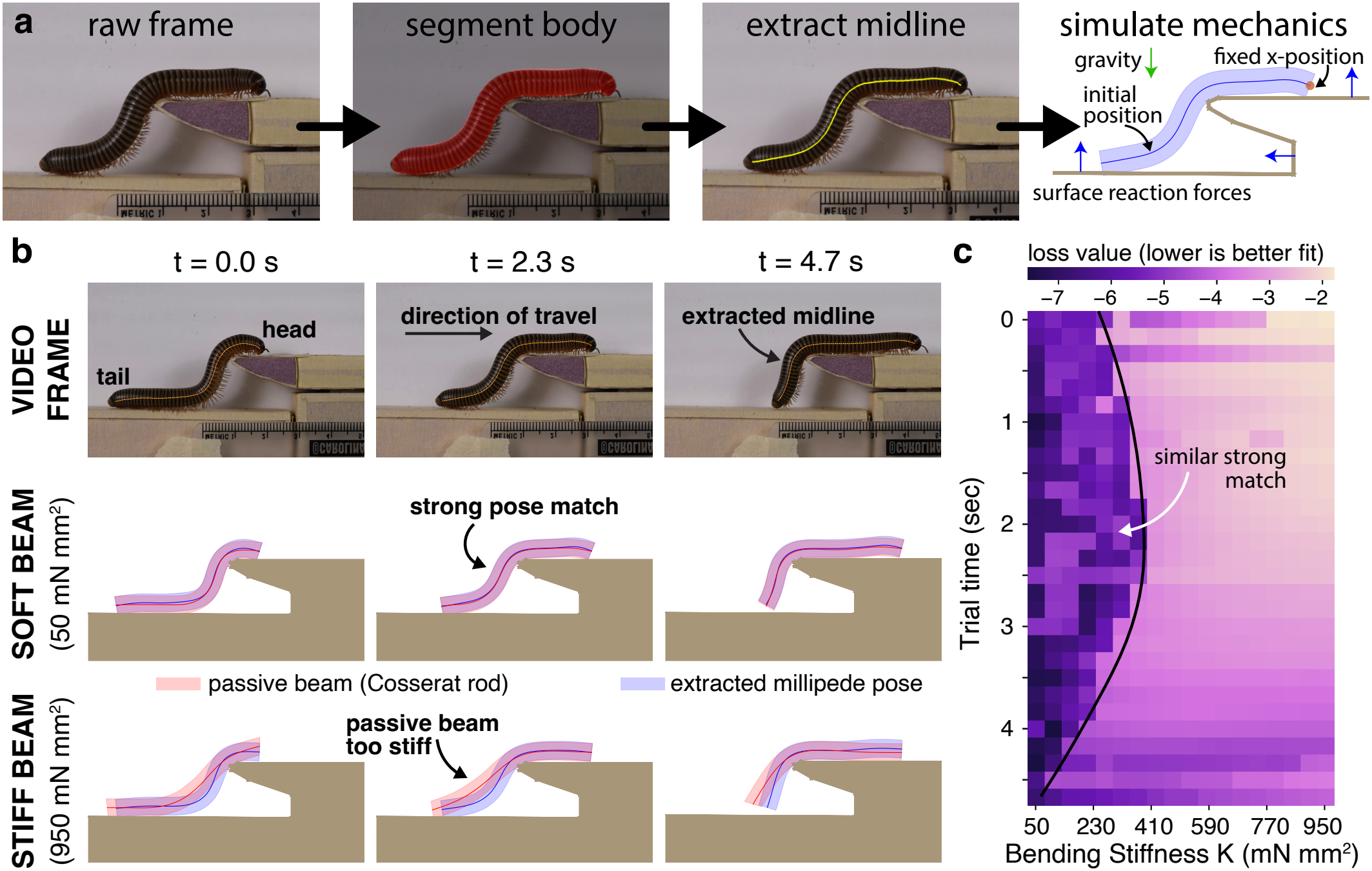
Modeling millipede trunk pose as a passive Cosserat rod. (a) Overview of segmentation and modeling pipeline. (b) Snapshots of millipede at different stages of traversing an upward gap and comparison of Cosserat rod passive rod simulations of different bending stiffnesses to the extracted millipede pose. In the bottom two rows, the blue shaded area is the extracted millipede pose. The red is the beam position after reaching equilibrium. In the middle row, a bending modulus of 50 mN·mm2, visually aligns with the millipede’s position while a bending modulus of 950·mN mm2 is straighter than the initial position. (c) The log mean square difference (loss) between the millipede’s pose and the equilibrium pose of a soft rod with a specified bending stiffness for each frame of the trial. Softer bending moduli generally produced the lowest loss and best match with the millipede’s pose across all frames.

For each video, millipede diameter and terrain locations were manually labeled and millipede midline tracking and ruler extraction were validated. Finally, the frames when the millipede first reached contact with a new terrain level and began crawling as well as when they lost contact with the beginning terrain were manually labeled to define a gap crossing interval.

Cosserat rod simulations. The millipede was simulated as a Cosserat rod, which is a slender, 1-D representation of an elastic filament capable of undergoing all modes of deformation [35, 36]. Although Cosserat rod formulations are capable of accounting for active muscle forces in slender animal biomechanics [36–41], since our goal here is to investigate if the millipede deviates from passive bending, we choose to model the millipede as a passive planar rod.

The exoskeleton of millipedes consists of overlapping rigid plates, while its body consists of ball-and-socket joints connecting tensile-force-resistant rings [3]. This morphology allows it to tightly curl up while resisting axial strain from anterior and posterior forces that arise during burrowing [18, 42]. This organization is not well-approximated by treating the millipede as a solid linear elastic rod where axial and bending stiffness are coupled through a single elastic (Young’s) modulus. Instead, we decouple the axial and bending stiffness of our rod. We fix the axial elastic modulus (*E*) of the millipede to be 20 kPa, which was found to be sufficient to limit any axial deformation, in line with millipedes’ general inextensibility [18]. We further fix the rod’s shear modulus (*G*) to be 100 kPa, which renders the rod effectively unshearable. Bending stiffness (*K*) is then set independently. While bending stiffness is often defined as *K = EI* where *I* is the second moment of area, here our decoupling of axial and bending stiffness allows us to directly define *K*.

We implement our model using *Elastica* [43], an open-source Cosserat-rod framework in Python that has been applied to and validated against a wide set of slender-body mechanics problems, including animal locomotion and manipulation [36–38, 41], continuum robotic control [44–46], and fibrous meta-materials [47, 48].

To analyze the deviation of the millipede from a passive rod, we adopt a quasi-static perspective. For every frame, a rod with a specified bending stiffness is initialized using the extracted millipede midline position and radius. The rod is then timestepped forward to allow it to relax its shape until equilibrium, defined as the change in energy over a window of five time periods α each below a threshold ϵ. Here, α = 50 µs and ϵ = 20 nJ. This resulted in simulations typically taking between 0.2 and 16 seconds to reach equilibrium. To avoid slipping down the gap, the end of the rod higher in the y-direction was fixed in its x-direction but free to rotate and move in other directions. The millipede’s density was approximated as that of water (0.997 g/mL). Gravity was applied as an external force, and ground reaction forces between the millipede and the surfaces were implemented using a spring-damper response force in line with our previous work [38, 41].

If the initialized rod position was not compatible with a passive rod with the specified bending stiffness, the rod would deform into a new equilibrium shape. The difference between the initial (x0) and equilibrium (xe) shape was then quantified as the mean-square error along its length 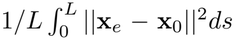 to define a loss. Minimizing this loss yields the passive bending stiffness that best matches the millipede pose of that frame. Because the segmentation process used to extract the midline resulted in jagged position and curvature values, the initial position x0 was defined as the rod’s position after 25 ms, which allows initial numerical jaggedness to rapidly smooth out without changing the overall position of the beam. For each frame, a parameter sweep of bending stiffness was performed over a range of *K*. The bending stiffness range was determined based on *K* = βπr4/4 where *r* is the extracted millipede radius and β ∈ [1, 18] kPa. This resulted in a typical range of bending stiffness between 20 and 1300 mN·mm2 which encompasses the range of biologically plausible bending stiffnesses with the lower bound on the order of ultra soft hydrogel (~50x softer than tofu [49]). At such low bending stiffnesses (and those lower), the beam acts effectively as an inextensible chain with no bending rigidity, capturing the physical lower bound of bending stiffness.

## RESULTS

Results from a representative case of a millipede traversing an upward step are shown in Fig. 1. Evaluating frames from the beginning, middle, and end of the trial show that in all cases the configuration of the millipede is best described by a rod with low bending stiffness (Fig. 1b). Overlaying the passive beam configuration (red) and the extracted millipede configuration (blue) for the two extremes of bending stiffness investigated (50 and 950 mN·mm2) shows that a softer rod provides a close match to the pose of the millipede while stiffer rods do not conform sufficiently, with their internal elastic stiffness straightening the millipede body across the gap.

The evolution of how rods of varying stiffness match the millipede throughout the trial (Fig. 1c) shows that, while 50 mN·mm2 is consistently the lowest loss value, there is a threshold bending stiffness of approximately 200-300 mN·mm2 below which decreasing bending stiffness does not substantially change in loss. At the end of the trial, there is decrease in loss values across all evaluated bending stiffness values. This corresponds to the millipede’s body having primarily completed its traverse of the step. At this point, most of the millipede’s trunk is on the top step in a straight configuration (such as at t = 4.7 s) and so is equally well described by any bending stiffness.

The combined results of all trials for traversing both upward and downward steps are shown in Fig. 2a-d. To compare across trials, trials are reported and aggregated in terms of their normalized time *t/W* where *W* is the time between when the millipede first makes contact with the top terrain, thus bridging the two terrain levels, to where the millipede leaves contact with the bottom terrain, thus becoming unsupported at the back. The millipede during this crawling-phase maintains a relatively consistent velocity throughout each video’s traversal window.

**FIG 2.**
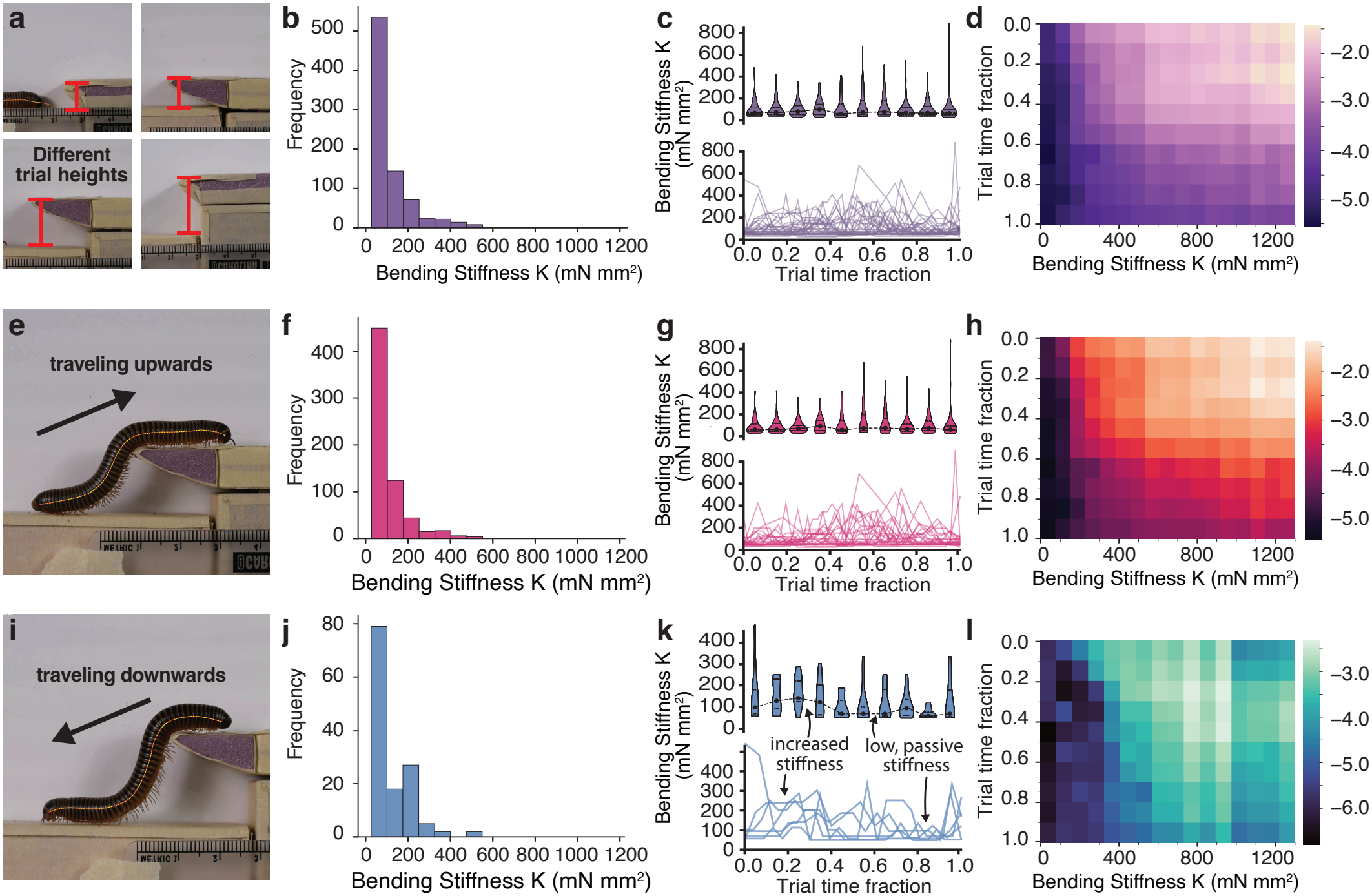
Aggregated simulation results over all trials. (a) Examples of the different heights and overhangs used in terrain bridging trials. (b) Histogram of the aggregated bending stiffnesses that yielded the lowest loss for each frame and for all trials showing softer bending stiffnesses, and so minimal stiffening along the trunk, consistently best describe the millipede pose for all cases considered. (c) Top: violin plot of distribution of bending stiffnesses with lowest loss for each frame and all trials showing evolution of the best matching bending stiffness over a terrain bridging behavior. Results are grouped by normalized trial time into decile groupings. Bottom: evolution of best matching bending stiffnesses for individual trials, showing no trials yielded consistently higher bending stiffness. (d) Heatmap of the log mean square error of the initial and steady-state positions normalized over the rod length for each frame aggregated into decile groupings and averaged together. (e–h) The same data as in (a–d) but broken out to show results of only upward trials. The upwards locomotion data aligns with the aggregated data. (i–l) The same data as in (a–d) but broken out to show results of only downward trials. The downwards data also skews towards a soft bending modulus, but also displays a smaller peak around 300 mN·mm2 possibly suggesting there may be some selective stiffening during downwards locomotion.

A histogram of the best bending moduli of all cases combined (Fig. 2b) illustrates how overall a very compliant, passive beam best describes the configuration of the millipede. Violin plots of the distribution of bending modulus that best match the millipede’s pose at each point of the traverse, as well as traces of the best bending modulus for each individual trial, further illustrate that across the entire traversal behavior, a low bending stiffness best describes millipede pose and stiffness. Temporary spikes of individual trials are transient and likely a result of multiple bending stiffnesses providing similar matching of the pose. Indeed, Fig. 2d shows that, similar to Fig. 1c, when aggregated over all trials, a similar plateau of similarly low loss values appears for bending stiffness lower than ~200 mN·mm2.

When broken out by traversing up a step (Fig. 2e-h) compared to down a step (Fig. 2i-l), low bending stiffness well describes the millipede’s position when traveling upward. When traveling down the step, while a low bending stiffness well describes the milipede stiffness, a second smaller peak appears at 300 mN·mm2 (Fig. 2j). This increased bending stiffness appears in the time distribution at 20–30% of the traversal behavior (Fig. 2k). This aligns with when the millipede has completed a bridge between the upper and lower step and begins to lower its legs and trunk to comply with the surface. The resulting increase in stiffening over this interval suggests the millipede may actively stiffen muscles in its trunk to stabilize its posture until more contact with the lower step is achieved, at which point the millipede returns to a passive, compliant bending stiffness. Alternatively, there may be anatomical differences in the most anterior segments which are not externally apparent, resulting in regional differences dorsal vs ventral bending stiffness.

## DISCUSSION AND CONCLUSION

Our results suggest the American giant millipede (*N. americanus*) does not display strong or regionalized muscle contraction to stabilize itself or otherwise modify its pose when its middle section is unsupported as it crosses a stepped terrain, nor does this stiffness vary substantially over time. Contrary to active regional bending, where regions of higher and lower bending stiffness would be expected, the millipede maintains a very low, uniform bending stiffness across its whole body.

This uniform stiffness could arise from low but uniform activation of the trunk muscles, pressurization of the haemocoelom cavity, or simply the animals passive body stiffness. Distinguishing between these possibilities is beyond the scope of this paper. However, based on the extremely low bending stiffness values found to match the millipede pose, active stiffening would incur a metabolic cost for unclear mechanical or stability benefit [50]. As such, it seems instead the millipede is using its body’s passive compliance without active stabilization when its middle section is unsupported during terrain bridging.

Our results generally find that the lowest bending stiffness best matches the millipede’s trunk pose. This naturally poses the question of if lower bending stiffness might provide even better tracking of the trunk’s pose. However, as described in the methods, our lower bound already corresponds to the fundamental lower limit of a beam acting effectively as an inextensible chain of links. Further, for the low bending stiffnesses cases where there is a close match between the millipede and the model’s pose (e.g., Fig. 1c), minute local discrepancies come to dominate the loss function as there is strong agreement for all low bending cases.

This use of body compliance may be viewed as an example of mechanically intelligent behavior [13, 14] by the millipede, as it provides a simple yet robust method for crossing heterogeneous, uneven terrain that requires minimal or no neuromuscular control. This is in contrast to the behavior of various snakes, which also have slender, compliant bodies, but who actively use muscles along their trunk to stabilize themselves during similar terrain bridging [9].

Millipedes did actively manipulate their trunks to initially bend their body upwards or stabilize their trunk as they moved downward when their head was unsupported as it worked to initially bridge the step. Our modeling framework could, in the future, be extended to incorporate active millipede muscle forces to identify the active forces and couples developed during this behavior. This active behavior may also contribute to the small transient stiffening behavior seen for the millipede traveling down a step as the millipedes relaxes muscle co-contraction used to stabilize itself. Once establishing contact on both ends of its body, however, the millipede adopts a passive response. We note that the behavior a millipede might adopt when traversing a larger gap is unclear. There may be a critical minimum level of body support beyond which trunk stiffening is employed. However, such a scenario is also limited by gap sizes the millipede can initially cross when supported by only one side, which may be a stronger constraint.

Overall, our results indicate that the pose of *N. americanus* is highly consistent with a passive elastic rod draped across the gap and we find no evidence for active regional stiffening or bending as it crosses the gap. These results advance understanding of how passive, mechanical intelligence in compliant slender organisms facilitates robust locomotion in heterogeneous terrains. They also hold potential to aid design and control of future soft robots that incorporate these principles to increase functionality.

## Acknowledgments

Support for this work was provided by National Science Foundation awards CMMI–#2528850 and EFRI– #2422340 (N.N).

## Data Availability

This work utilized the open-source Elastica software package available online at https://github.com/GazzolaLab/PyElastica [43]. Code for the setup and simulation of the millipede stiffness tests performed in this work will be released on GitHub upon final publication.

## References

[1] S. A. Wainwright, Axis and circumference: the cylindrical shape of plants and animals. Harvard University Press, 1988.

[2] B. Chong, J. He, D. Soto, T. Wang, D. Irvine, G. Blekherman, and D. I. Goldman, “Multilegged matter transport: A framework for locomotion on noisy landscapes,” Science, vol. 380, no. 6644, pp. 509–515, 2023.

[3] S. M. Manton, “The evolution of arthropodan locomotory mechanisms.—part 4. the structure, habits and evolution of the diplopoda,” Zoological Journal of the Linnean Society, vol. 42, pp. 299–368, 1954.

[4] S. P. Hopkin and H. J. Read, The biology of millipedes. Oxford University Press, 1992.

[5] H. C. Astley, “Long limbless locomotors over land: the mechanics and biology of elongate, limbless vertebrate locomotion,” Integrative and Comparative Biology, vol. 60, no. 1, pp. 134–139, 2020.

[6] W. M. Kier and K. K. Smith, “Tongues, tentacles and trunks: the biomechanics of movement in muscular-hydrostats,” Zoological Journal of the Linnean Society, vol. 83, no. 4, pp. 307–324, 1985.

[7] W. M. Kier, “The diversity of hydrostatic skeletons,” Journal of Experimental Biology, vol. 215, no. 8, pp. 1247–1257, 2012.

[8] N. Naughton, “Muscular hydrostats,” Current Biology, vol. 36, no. 19, pp. R1012–R1016, 2026.

[9] R. M. Jorgensen and B. C. Jayne, “Three-dimensional trajectories affect the epaxial muscle activity of arboreal snakes crossing gaps,” Journal of Experimental Biology, vol. 220, no. 19, pp. 3545–3555, 2017.

[10] B. C. Jayne, “Muscular mechanisms of snake locomotion: an electromyographic study of lateral undulation of the florida banded water snake (nerodia fasciata) and the yellow rat snake (elaphe obsoleta),” Journal of Morphology, vol. 197, no. 2, pp. 159–181, 1988.

[11] C. O. Bennice, K. C. Buresch, J. H. Grossman, T. D. Morano, and R. T. Hanlon, “Octopus arm flexibility facilitates complex behaviors in diverse natural environments,” Scientific Reports, vol. 15, no. 1, p. 31875, 2025.

[12] T. Flash and L. Zullo, “Biomechanics, motor control and dynamic models of the soft limbs of the octopus and other cephalopods,” Journal of Experimental Biology, vol. 226, Apr. 2023.

[13] T. Wang, C. Pierce, V. Kojouharov, B. Chong, K. Diaz, H. Lu, and D. I. Goldman, “Mechanical intelligence simplifies control in terrestrial limbless locomotion,” Science Robotics, vol. 8, no. 85, p. eadi2243, 2023.

[14] P. E. Schiebel, J. M. Rieser, A. M. Hubbard, L. Chen, D. Z. Rocklin, and D. I. Goldman, “Mechanical diffraction reveals the role of passive dynamics in a slithering snake,” Proceedings of the National Academy of Sciences, vol. 116, no. 11, pp. 4798–4803, 2019.

[15] P. E. Marek and W. A. Shear, “Myriapods,” Current Biology, vol. 32, p. R1294–R1296, Dec. 2022.

[16] A. Garcia, S. Priya, and P. Marek, “Understanding the locomotion and dynamic controls for millipedes: part 1—kinematic analysis of millipede movements,” in ASME 2015 Conference on Smart Materials, Adaptive Structures and Intelligent Systems, vol. 57304, p. V002T06A005, American Society of Mechanical Engineers, 2015.

[17] H. M. Wilson, “Muscular anatomy of the millipede phyllogonostreptus nigrolabiatus (diplopoda: Spirostreptida) and its bearing on the millipede “thorax”,” Journal of Morphology, vol. 251, no. 3, pp. 256–275, 2002.

[18] A. Garcia, G. Krummel, and S. Priya, “Fundamental understanding of millipede morphology and locomotion dynamics,” Bioinspiration & Biomimetics, vol. 16, no. 2, p. 026003, 2021.

[19] J. A. Nirody, “Flexible locomotion in complex environments: the influence of species, speed and sensory feedback on panarthropod inter-leg coordination,” Journal of Experimental Biology, vol. 226, no. Suppl_1, p. jeb245111, 2023.

[20] T. Kano, K. Sakai, K. Yasui, D. Owaki, and A. Ishiguro, “Decentralized control mechanism underlying interlimb coordination of millipedes,” Bioinspiration & Biomimetics, vol. 12, no. 3, p. 036007, 2017.

[21] S. Manton, “The evolution of arthropodan locomotory mechanisms—part 3. the locomotion of the chilopoda and pauropoda.,” Zoological Journal of the Linnean Society, vol. 42, no. 284, pp. 118–167, 1952.

[22] J. Hou, Z. Xue, Y. Liang, Y. Sun, Y. Zhao, and Q. Chen, “Self-configurable centipede-inspired rescue robot,” Applied Sciences, vol. 14, no. 6, p. 2331, 2024.

[23] S. J. Saha, M. A. Chowdhury, S. Islam, S. A. Deowan, S. E. Arman, and A. K. Ghosh, “Flexipede: A bio-inspired, modular myriapod robot for rough-terrain traversal,” Robotics, vol. 15, no. 7, p. 129, 2026.

[24] K. Yasui, A. Ohno, T. Kano, and A. Ishiguro, “Decentralized control law for load-adaptive gaits of multi-legged robots inspired by millipedes,” Advanced Robotics, vol. 39, no. 1, pp. 79–88, 2025. doi: 10.1080/01691864.2024.2408619.

[25] H. Yin, R. Shi, and J. Liu, “Structural design and control research of multi-segmented biomimetic millipede robot,” Biomimetics, vol. 9, no. 5, p. 288, 2024.

[26] K. Yasui, T. Yamaichi, and A. Ishiguro, “Millipedes suggest decentralized trunk–limb coordination control is essential for inter-limb coordination in many-legged locomotion,” Artificial Life and Robotics, vol. 31, no. 1, pp. 34–45, 2026.

[27] K. Diaz, E. Erickson, B. Chong, D. Soto, and D. I. Goldman, “Active and passive mechanics for rugose terrain traversal in centipedes,” Journal of Experimental Biology, vol. 226, no. 4, p. jeb244688, 2023.

[28] K. Yasui, S. Takano, T. Kano, and A. Ishiguro, “Adaptive centipede walking via synergetic coupling between decentralized control and flexible body dynamics,” Frontiers in Robotics and AI, vol. 9, p. 797566, 2022.

[29] B. D. Anderson, J. W. Shultz, and B. C. Jayne, “Axial kinematics and muscle activity during terrestrial locomotion of the centipede Scolopendra heros,” Journal of Experimental Biology, vol. 198, no. 5, pp. 1185–1195, 1995.

[30] C. Liu, Z. You, and P. Maiolino, “Kinematics of an origami inspired millipede robot,” in International Design Engineering Technical Conferences and Computers and Information in Engineering Conference, vol. 86281, p. V007T07A062, American Society of Mechanical Engineers, 2022.

[31] R. Khanam and M. Hussain, “YOLOv11: An overview of the key architectural enhancements,” arXiv preprint arXiv:2410.17725, 2024.

[32] G. Jocher and J. Qiu, “Ultralytics YOLO11,” 2024. Version 11.0.0; AGPL-3.0.

[33] B. Dwyer, J. Nelson, T. Hansen, et al., “Roboflow (version 1.0),” 2024. Computer vision software.

[34] R. O. Duda and P. E. Hart, “Use of the Hough transformation to detect lines and curves in pictures,” Communications of the ACM, vol. 15, no. 1, pp. 11–15, 1972.

[35] S. S. Antman, “Problems in nonlinear elasticity,” Nonlinear Problems of Elasticity, pp. 513–584, 2005.

[36] M. Gazzola, L. H. Dudte, A. G. McCormick, and L. Mahadevan, “Forward and inverse problems in the mechanics of soft filaments,” Royal Society Open Science, vol. 5, no. 6, 2018.

[37] X. Zhang, F. K. Chan, T. Parthasarathy, and M. Gazzola, “Modeling and simulation of complex dynamic musculoskeletal architectures,” Nature communications, vol. 10, no. 1, pp. 1–12, 2019.

[38] X. Zhang, N. Naughton, T. Parthasarathy, and M. Gazzola, “Friction modulation in limbless, three-dimensional gaits and heterogeneous terrains,” Nature communications, vol. 12, no. 1, p. 6076, 2021.

[39] H.-S. Chang, U. Halder, C.-H. Shih, N. Naughton, M. Gazzola, and P. G. Mehta, “Energy-shaping control of a muscular octopus arm moving in three dimensions,” Proceedings of the Royal Society A, vol. 479, no. 2270, p. 20220593, 2023.

[40] C.-H. Shih, N. Naughton, U. Halder, H.-S. Chang, S. H. Kim, R. Gillette, P. G. Mehta, and M. Gazzola, “Hierarchical control and learning of a foraging cyberoctopus,” Advanced Intelligent Systems, vol. 5, no. 9, p. 2300088, 2023.

[41] A. Tekinalp, N. Naughton, S. H. Kim, U. Halder, R. Gillette, P. G. Mehta, W. Kier, and M. Gazzola, “Topology, dynamics, and control of a muscle-architected soft arm,” Proceedings of the National Academy of Sciences, vol. 121, no. 41, p. e2318769121, 2024.

[42] B. J. Borrell, “Mechanical properties of calcified exoskeleton from the neotropical millipede, nyssodesmus python,” Journal of Insect Physiology, vol. 50, no. 12, pp. 1121–1126, 2004.

[43] A. Tekinalp, S. H. Kim, Y. Bhosale, T. Parthasarathy, N. Naughton, A. Albazroun, R. Joon, S. Cui, I. Nasiriziba, M. Stölzle, C.-H. C. Shih, and M. Gazzola, “Gazzolalab/pyelastica,” 2024.

[44] N. Naughton, J. Sun, A. Tekinalp, T. Parthasarathy, G. Chowdhary, and M. Gazzola, “Elastica: A compliant mechanics environment for soft robotic control,” IEEE Robotics and Automation Letters, vol. 6, no. 2, pp. 3389–3396, 2021.

[45] N. Naughton, A. Tekinalp, K. Shivam, S. H. Kim, A. Khairnar, V. Kindratenko, and M. Gazzola, “Neural reservoir control of a bio-hybrid soft arm,” Proceedings of the National Academy of Sciences, vol. 123, no. 17, p. e2522094123, 2026.

[46] A. Boron, B. Zheng, Z. Zhou, N. Naughton, and S. Li, “Stochastic entanglement of deterministic origami tentacles for robust robotic grasping,” Advanced Science, p. e76810, 2026.

[47] Y. Bhosale, N. Weiner, A. Butler, S. H. Kim, M. Gazzola, and H. King, “Micromechanical origin of plasticity and hysteresis in nestlike packings,” Physical review letters, vol. 128, no. 19, p. 198003, 2022.

[48] A. Khairnar, Y. Phalak, J. Wang, Z. Zhou, B. Jantzen, S. Li, and N. Naughton, “Spider web-inspired sensing and computation with fiber network physical reservoirs,” Journal of Intelligent Material Systems and Structures, p. 1045389X261470871, 2026.

[49] Institute of Physics, “Young’s (or should it be yum’s?) modulus of food and other measurements.” IOP SPARK, Stories from Physics: Properties of Matter, n.d. Accessed: 2026-11-09.

[50] N. Hogan, “Adaptive control of mechanical impedance by coactivation of antagonist muscles,” IEEE Transactions on automatic control, vol. 29, no. 8, pp. 681–690, 1984.

